# The Hidden Biocatalytic Potential of the Old Yellow Enzyme Family

**DOI:** 10.1101/2023.07.10.548207

**Authors:** David W. White, Samantha Iamurri, Parisa Keshavarz-Joud, Tamra Blue, Janine Copp, Stefan Lutz

## Abstract

The rapid advancement of sequencing technology has created an immense reservoir of protein sequence-function information that has yet to be fully utilized for fundamental or biocatalytic applications. For example, ene reductases from the ‘old yellow enzyme’ (OYE) family catalyze the asymmetric hydrogenation of activated alkenes with enhanced stereoselectivity - key transformations for sustainable production of pharmaceutical and industrial synthons. Despite the proven biocatalytic application, the OYE family remains relatively underexplored with only 0.1% of identified members having any experimental characterization. Here, a platform of integrated bioinformatics and synthetic biology techniques was employed to systematically organize and screen the natural diversity of the OYE family. Using protein similarity networks, the known and unknown regions of the >115,000 members of the OYE family were broadly explored to identify phylogenetic and sequence-based trends. From this analysis, 118 novel enzymes were characterized across the family to broadly explore and expand the biocatalytic performance and substrate scope of known OYEs. Over a dozen novel enzymes were identified exhibiting enhanced catalytic activity or altered stereospecificity. Beyond well-established ene reduction, we detected widespread occurrence of oxidative chemistry amongst OYE family members at ambient conditions. Crystallography studies of selected OYEs yielded structures for two enzymes, contributing to a better understanding of their unique performance. Their structures revealed an unusual loop conformation within a novel OYE subclass. Overall, our study significantly expands the known functional and chemical diversity of OYEs while identifying superior biocatalysts for asymmetric reduction and oxidation.

## Introduction

Over the past two decades biocatalysis has fundamentally changed chemical production by utilizing the high selectivity of enzymes to create more streamlined and sustainable processes.^1–5^ However, enzymes are not designed for industrial processes as they suffer from compromised stability and limited substrate scope.^6, 7^ Protein engineering through directed evolution or computational redesign are well-proven strategies in developing enzymes for biocatalysis, but remains a relatively slow, low-yielding, and labor-intensive process.^7, 8^ The discovery of novel biocatalysts could hasten this process and help meet the growing demand for biocatalysis in industry.

The rapid development of sequencing technology has revealed a near-limitless reservoir of potential biocatalysts in the millions of uncharacterized protein sequences that await in ever-growing databases.^9–11^ Although uncharacterized, these sequences have evolved towards a specific function which may be useful in biotransformations. Exploration of this vast uncharacterized sequence space through enzyme family profiling has become a promising approach for novel biocatalyst identification with studies investigating halogenases, phosphatases, methyltransferases, dehalogenases, and imine reductases.^12–16^ The natural diversity of enzymes capable of asymmetric hydrogenation, a key reaction for chiral synthesis, has yet to be evaluated.

Old Yellow Enzymes (OYEs) (Fig. 1a) offer a green alternative to known chiral transition metal complexes and organocatalysts that catalyze asymmetric trans-hydrogenation of activated alkenes with chemo-, regio- and stereo-selectivity (Fig. 1b). Consequently, OYEs have been employed for the production of diverse products, including anti-inflammatory drugs, macrocyclic antibiotics, and anti-convulsants. OYE family enzymes are TIM barrel fold flavoproteins that bind and utilize a flavin mononucleotide cofactor for hydride transfer to activated alkenes. Previous studies have delineated the OYE family into five classes (Class I, II, III, IV, and V) based on the conservation of specific residues, structure, and phylogeny.^17^ Since their discovery in 1932, only 0.1% of OYEs have any experimental characterization.^18^ Furthermore, the lack of a native substrate for most enzymes combined with the newly identified classes of OYEs hint at genetic and catalytic diversity of the family.^17, 19^ With over 115,000 uncharacterized sequences, it is likely that the full biocatalytic potential of OYEs has yet to be unlocked.

**Figure 1.**
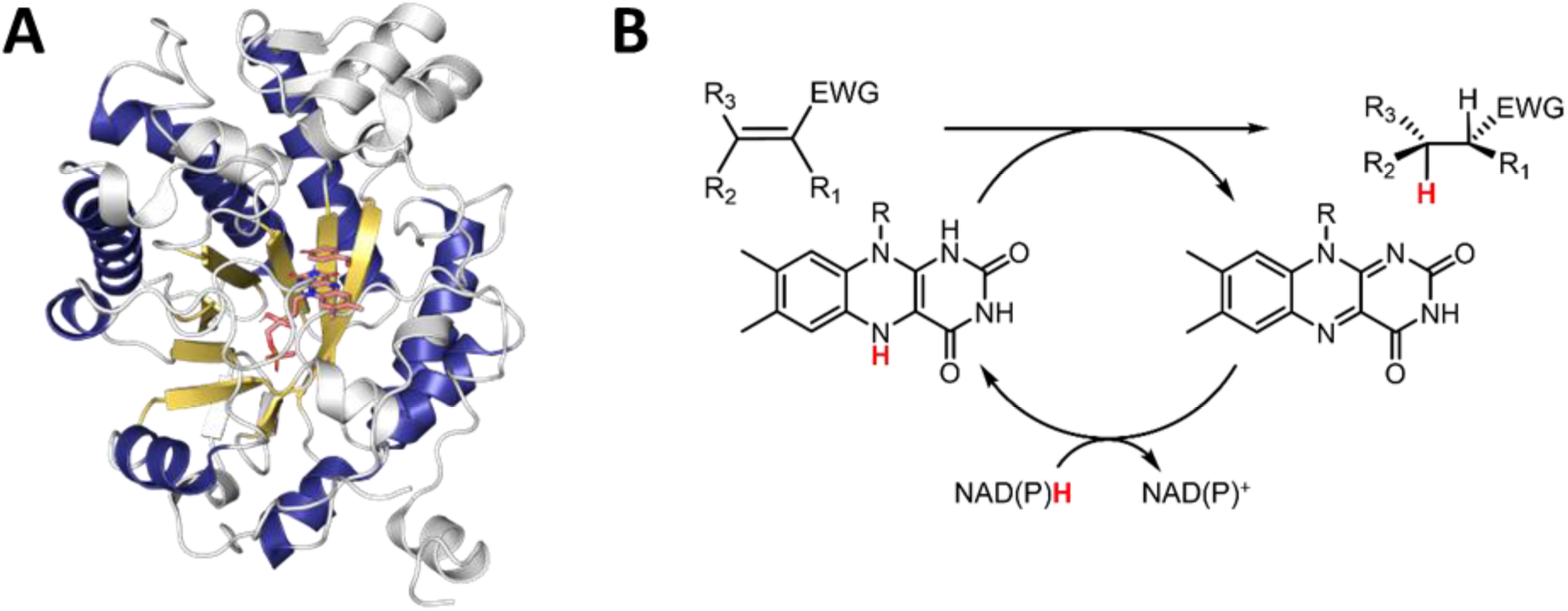
Generalized OYE structure and mechanism. A) The TIM barrel fold of OYEs with the repeating α-helix (dark blue) and β-sheets (gold) highlighted. The FMN and inhibitor (orange) are also present. (PDB: 1OYB) B) The catalytic cycle of OYEs for asymmetric hydrogenation. A hydride (red) is transferred from NAD(P)H to the FMN cofactor. The hydride is transferred to the β of the α,β-unsaturated compound.

Here we describe the systematic organization and screening of the diverse OYE family using bioinformatic and synthetic biology techniques to explore the biocatalytic potential hidden within its uncharacterized sequence space.

## Results and Discussion

### Bioinformatic-based Organization of the Old Yellow Enzyme Family

A comprehensive overview of the OYE family was constructed using an SSN of 70,366 OYE sequences at the edge threshold of 10^-85^ (Fig. 2A). This threshold was selected as it organized the known OYE classes into distinct sequence clusters. At this threshold, sequences with greater than ∼35% sequence identity group together with many clusters containing enzymes with specific conserved residues or motifs. Overall, the majority (∼85%) of OYE sequences are bacterial in origin and range between 300-450 amino acids in length - though enzymes were found in all kingdoms of life and in various sizes. The SSN analysis resulted in 20 major clusters, each containing >100 unique sequences and combined, representing 99.8% of the sequences present in the SSN. Of the major clusters, 10 contained at least one of the 126 previously reported OYEs (Fig. 2A), however, only 5 of these clusters were represented by more than one characterized enzyme. The other 10 clusters have no known biological roles or documented activities.

**Figure 2.**
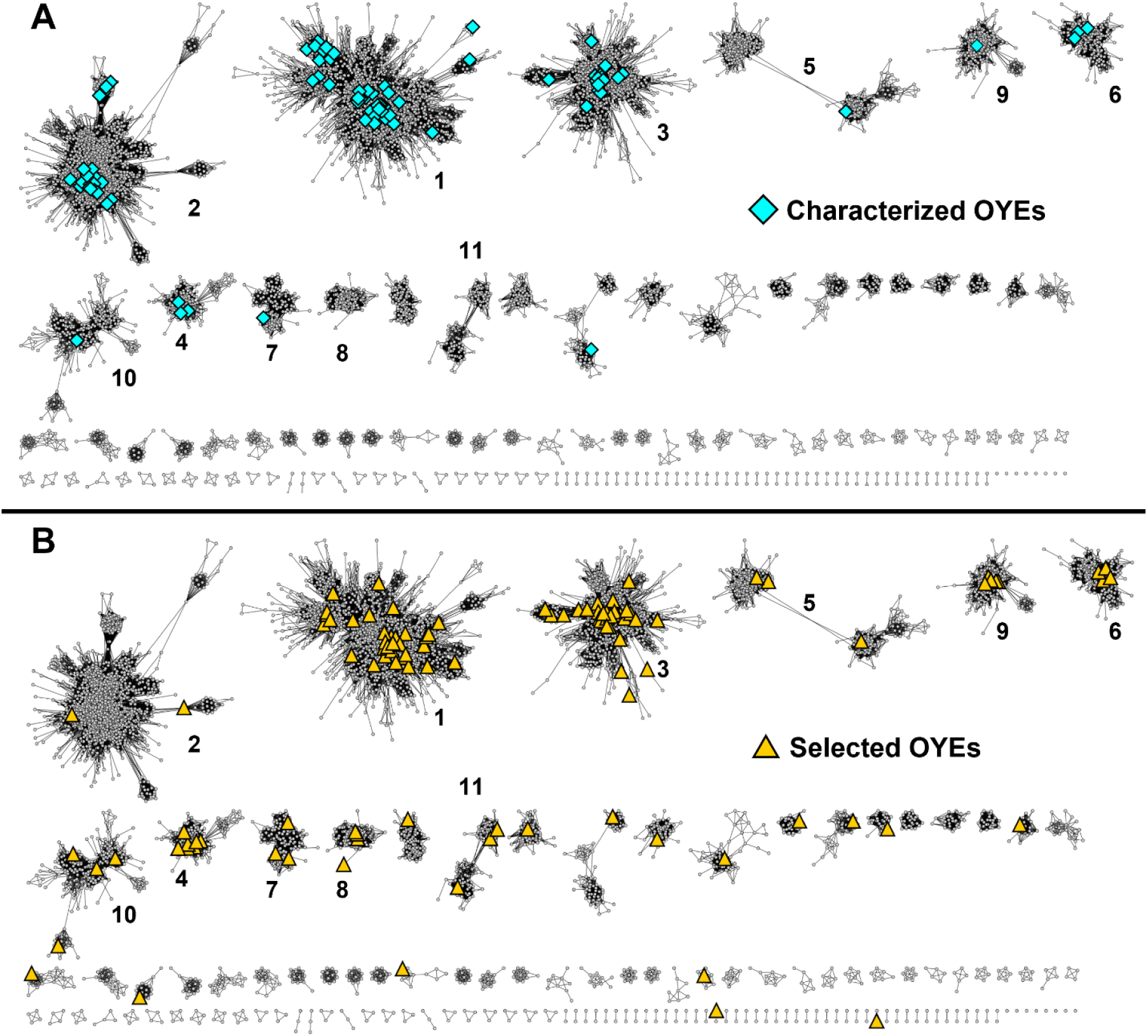
Two views of the SSN of the OYE family. Numbering is based on the size or number of sequences in the cluster (i.e. cluster 1 is the largest). A) The previously characterized OYEs (blue diamonds) are highlighted in the SSN map showing the explored clusters. B) The same view of the OYE family but displaying the screened sequences (orange triangles) in this study. 50% ID for nodes and E-value threshold of 10^-85^. Created in Cytoscape.

### Individual Cluster Analysish

Clusters 1 through 3 represent the three largest groups, each containing ∼25% of OYE family members. The largest group, cluster 1, comprises the previous classification of Class I or ‘classical’ OYEs, including the fungal OYE1-like, the plant OPR-like, and the bacterial PETNR- like enzymes.^19^ Half of all characterized OYEs are located within this group, including the majority of those with solved crystal structures.^19, 20^ Cluster 2 consists of the 2-enoate reductases, with the majority of these sequences containing the second FAD binding domain. The third largest group, cluster 3, encompasses the Class II or ‘thermophilic OYEs’.^21^ Two defining features of this class and observed in the cluster is the conserved cysteine residue (other OYEs contain a threonine) known to interact with the flavin cofactor as a redox modulator and the C-terminal ‘arginine finger’ that interacts with another monomer’s active site.^22^ Surprisingly, a significant portion (15%) of cluster 3 are the multidomain 2-aminobenzyoyl-CoA monooxygenase/reductases – a related OYE subgroup rarely associated with the family.^23^

The remaining clusters comprise roughly the last 25% of the OYE family. Unlike the three large clusters, the remaining groups are only lightly sampled or completely unexplored. Cluster 4 contains the characterized Class IV OYEs: Chr-OYE1, Rer-ER7, and Ppo-ER3. Enzymes from this cluster contain a ∼20 amino acid insertion in between β-sheet 7 and α-helix 7, in agreeance with previous analysis of Class IV enzymes.^17^ At the chosen SSN threshold, cluster 5 is composed of two loosely connected smaller subclusters contained entirely of either bacterial or fungal sequences. Within the fungal subcluster is the sole characterized cluster 5 OYE, IccE from *Penicillium variabile*.^24^ Cluster 6 consists primarily of Class III OYEs such as YqiG, LacER, and LrFOR, while cluster 13 contains the remaining Class III OYE: Lla-ER.^17^ Interestingly, nearly ∼20% of cluster 6 are >900 amino acid in sequence length with domains related to fumarate reductase/flavocytochrome C with one such enzyme already characterized.^25^

The sole representative of cluster 7 is Nox from *Rhodococcus erythropolis* MI2, an ‘orphan’ OYE that has yet to be organized within a class.^26^ Cluster 7 enzymes, including Nox, contain a conserved CxxCxxCx_n_C motif of a 4Fe-4S cluster near the C-terminus and aligns with the iron-sulfur cluster coordinating residues of 2-enoate reductases crystal structures. It is unclear what the function of the possible iron-sulfur cluster is for cluster 7 enzymes - whether structural or for electron transfer to/from an auxiliary binding partner. Previous characterization demonstrated that Nox could oxidize NADH, but it is unclear if a fully formed metal cluster was present in the enzyme during the assay.^26^ Regardless, this iron-sulfur cluster motif is also conserved in clusters 10, 14, 18, and 20.

The sole representative of Class V OYEs, ArOYE6 from *Ascochyta rabiei*, is in cluster 9. Recently, ArOYE6 was crystallized, revealing a novel C terminus region of ∼75 amino acids that was confirmed to be vital for FMN binding.^27^ Other sequences from this cluster also contain a similar C-terminus region that is typically 50-100 amino acids longer than canonical OYEs such as OYE1 and YqjM. Cluster 9 is also unique in that it is comprised solely of eukaryotic OYEs of which >20% originate from nematodes. Clusters 8, 11, 12, and 14-20 have yet to be characterized. Clusters beyond 13 begin to narrow in taxonomy and sequence similarity, likely indicating that these clusters may be small offshoots of larger clusters.

### Phylogenetic Analysis of the OYE Family

To investigate the relationships between the identified clusters, a phylogeny of the OYE family was generated using ∼500 representative sequences (Fig. 3). Enzymes from clusters 1-3 form distinct groups. Sequences from clusters 4 and 8 group together, as do those from clusters 6, 11, and 13, and 5, 7, 9, and 10. Cluster 12 does not group with other clusters.

**Figure 3:**
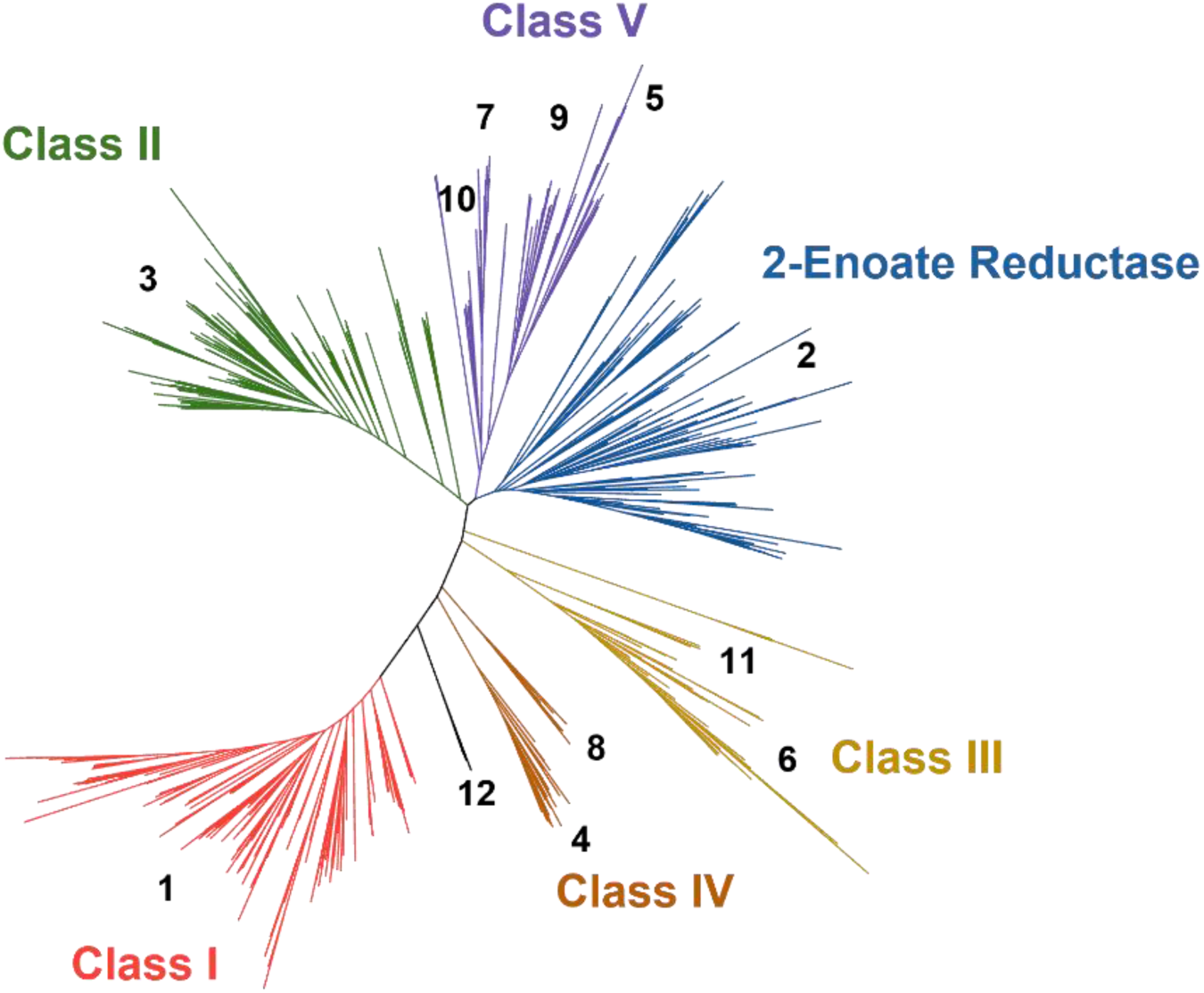
Unrooted phylogenetic tree of the OYE family. The approximate location of each cluster is noted in black. The updated classes are labeled using the previous colors of red for Class I, green for Class II, yellow for Class III, orange for Class IV, purple for class V, and blue for 2-Enoated reductases. The phylogenetic tree was formed using multiple sequence alignments using PROMALS3D, and the tree was calculated using IQ-Tree. Visualized using iTOL.

With further discovery of new OYE sequences, it is likely that these clusters will continue to change over time. To address the numerous attempts at classifying the OYE family, we propose the incorporation of these bioinformatic results into 5 OYE classes previously described by Peters et al (Fig. 3).^17^ To simplify, these 5 classes are kept with the addition of cluster designations for the unique clusters identified in this study. The revised classes are as follows: Class I: cluster 1, Class II: cluster 3, Class III: clusters 6, 11, and 13, Class IV: cluster 4 and 8, and Class V: clusters 5, 7, 9, and 10. The 2-enoate reductases of cluster 2 are excluded from this classification as these enzymes are historically considered a separate group.^28^ This cluster designation will be used for the remainder of this study.

### Selection of Novel OYE Sequences for In Vitro Screening

From among members of the SSN, 125 sequences were selected to systematically sample the OYE family (Fig. 2b; Table S1). This selection included 119 novel protein sequences and 6 known OYEs to serve as internal controls. Enzyme selection was guided by complementarity to known OYEs, proportional representation of cluster size, and variation in conserved active site regions (ex: T37, Y82, W116, and N194 in OYE1 numbering). Representatives from uncharacterized (sub)clusters and taxonomy were prioritized whilst avoiding candidates from organisms that may be problematic for protein expression in *E. coli*. Sequences of atypical length, < 240 aa and >555 aa, were avoided to eliminate protein fragments and multi-domain proteins that may be insoluble due to complex folding and unmet cofactor requirements. By these criteria, most enzymes from cluster 2 were eliminated from experimental consideration as they are typically >600 amino acids in length and contain multiple domains.

### Selection of Substrate Mixes, Reaction Conditions, and Characterized OYE Activity

For rapid, semi-quantitative functional screening, the selected sequences were synthesized using an in vitro transcription/translation system (IVTT).^29^ These reactions were supplemented with purified GroEL/GroES chaperones, FMN and FAD for proper folding and cofactor loading. As validation, IVTT screening results were confirmed with traditionally purified enzymes for novel OYEs. Likewise, solubility data for all library members were obtained in parallel by small-scale heterologous expression in *E. coli*. These studies suggest that 118 OYEs (95%) could be overexpressed; 87 (70%) produced a soluble fraction. Interestingly, protein insolubility did not correlate with the kingdom, affecting ∼30% of representatives from bacteria and fungi and ∼50% from archaea, plantae, metazoan and metagenomic DNA, respectively.

The selected OYEs were screened against 16 different substrates organized into 4 mixes (Fig. 4). Mix I contained standard OYE substrates, (*S*)-carvone (**1**), ketoisophorone (**2**), cinnamaldehyde (**3**), and 4-phenylbut-3-yn-2-one (**4**), which served as controls for typical ene reductase activity.^30–32^ Mix II consisted of various furanones (**5-8**) as precursors for high-value chiral synthons.^33^ Mix III represented challenging substrates that known OYEs have little to no activity for. The precursor to the Roche ester (**9**) is involved in various industrial synthesis and selective reduction of unsaturated nitriles (**10-11**) would be beneficial for ‘click’ chemistry.^34, 35^ Lastly, the nitro alkene (**12**) was incorporated to probe for nitro versus alkene reduction.^36^

**Figure 4:**
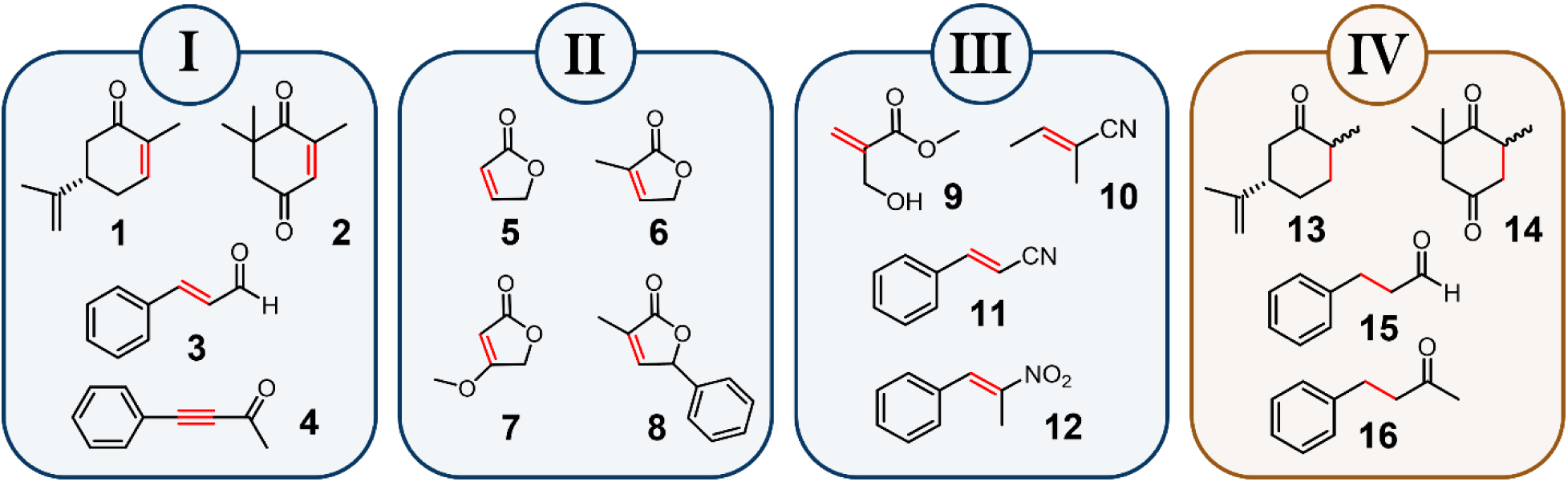
The substrate mixes. The 16 substrates used in this study are shown organized into 4 mixes. The color of the mix is the type of chemistry expected with blue for ene reduction and gold for desaturation. Highlighted in red is the bond which will be reduced (I-III) or desaturated (IV).

In addition to alkene reduction, growing interest in the desaturase activity of OYEs prompted the testing of reverse, oxidative chemistry for the selected sequences.^37^ Accordingly, mix IV used alkanes (**13-16**), the products of our four reference compounds, as substrates. Reactions with mix IV were conducted without nicotinamide cofactors and in the presence of atmospheric oxygen. Although reports have previously described OYE desaturase activity, it has required either artificial flavin cofactors or non-physiological temperatures.^38, 39^

The screening protocol was validated with six known OYEs (OYE1, NemA, YqiG, PETNR, FgaOx_3_, and TcOYE) to confirm the previously reported substrate conversion profiles.^40–43^ Of note, the closely related NemA and PETNR exhibited desaturase activity with mix IV and the expected conversion of substrates in mixes I and III. Additionally, this study significantly expanded the substrate scope of FgaOx_3_ from *Aspergillus fumigatus* and TcOYE from *Trypanosoma cruzi* – two enzymes previously studied only for native function.^42, 43^ FgaOx_3_ converted substrates in mixes I and III, while TcOYE displayed broad activity across all four reaction mixes and was the most promiscuous enzyme converting 9 of the 16 substrates.

### Overall OYE Activity and Cluster-Specific Trends

Of the novel 119 OYEs screened, 75 (60%) enzymes displayed activity for at least one substrate, representing 14 different clusters (Fig. 5). Activity was detected for 13 substrates; any enzyme did not detect a reduction of 7, 10, and 11. Alkene reduction of **12** was observed, but no nitro reduction was detected by any of the enzymes. Clear activity trends were recognized from specific clusters. For instance, cluster 1 enzymes have a variety of substrate profiles but overall preferred **12**. Likewise, cluster 3 showed preference for **1**, but could not readily convert **4** or **12**. This preference could be due to the geometry of the active site as cluster 1 enzymes conserve hydrophobic residues at the binding position of the phenyl group of **12** in the active site. In contrast, cluster 3 enzymes conserve the ‘arginine finger’ near this position in this active site, replacing this hydrophobic region with that of a charged residue.^20^ In other clusters, sequences from cluster 4 convert all of mix I while being inactive in mix II. Cluster 6 OYEs preferred to convert substrates **3** and **12** over other substrates – similar to cluster 1 enzymes. From the sequence info, it is unknown why clusters 4 and 6 display specific activity profiles.

**Figure 5:**
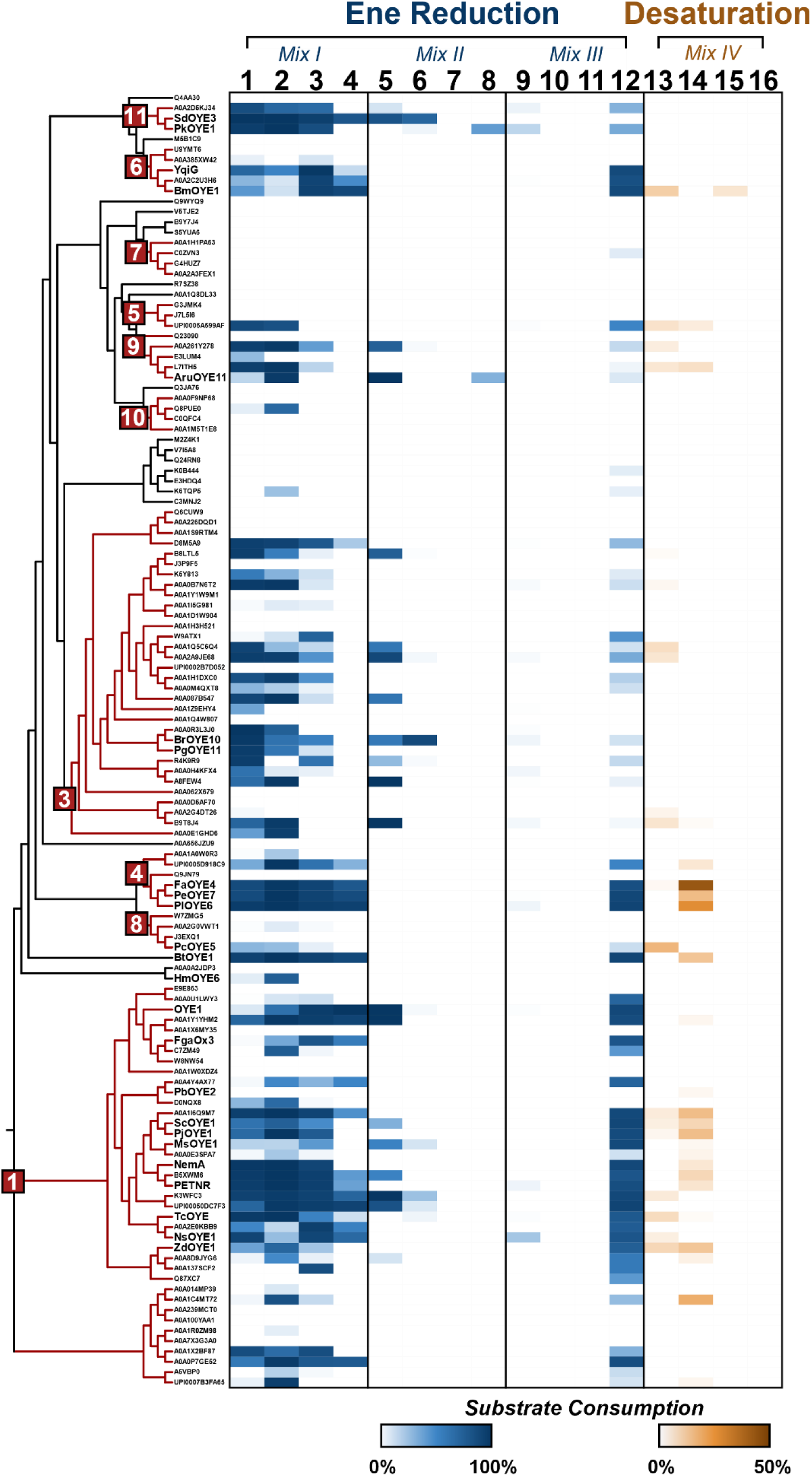
Heatmap of substrate consumption by the selected OYEs. The heatmap displays the percentage of substrate the selected sequences consumed in 24 hours. The darker the color, the more substrate consumed (blue for ene reduction, gold for desaturation). Desaturase activity is to 50% to provide help visualize the widespread activity. The substrates are labeled with boxes around the mixes. On the left side is a phylogenetic tree purely for organizing the sequences into clusters (red boxes and lines). Mentioned OYEs are named and uniprot codes are used for the remainder sequences. Created in iTOL.

Intriguingly, significant desaturase activity was observed across the family. Of the 125 enzymes screened, 33 enzymes from 8 distinct clusters converted at least one substrate in mix IV, indicating widespread desaturase activity within the OYE family (Fig. 5). Activity was commonly detected for substrates **13** and **14**, with enzymes and clusters typically showing a preference. For example, OYEs with desaturase activity in cluster 3 only convert **13,** while enzymes from cluster 4 show a strong preference for **14**. (Fig 5) This is similar to trends observed in the same clusters for the forward reaction. Notably, conversion for **15** was only detected by BmOYE1 from *Bacillus marmarensis*, an OYE from cluster 6, while no activity was observed for **16**.

### Interesting Biocatalytic Candidates of Mixes I-III

Several selected sequences displayed significantly improved or novel catalytic activity when compared to characterized enzymes such as OYE1 (Fig. 6A-G). In mix I, 3 enzymes reduced all substrates >85% within 24 hours (Fig. 6A). OYE1 in comparison reduces all mix I substrates but only converts <10% of **1** and ∼60% of **2**. Two enzymes are from cluster 4: PlOYE6 and PeOYE7 from *Pseudomonas lutea* and *Priestia endophytica* respectively, while the other OYE, BtOYE1 from *Bacillus thuringiensis,* is from the uncharacterized cluster 12. Mix 1 also probed enantioselectivity of the enzymes with substrates **1** and **2.** Notably, the majority of OYEs screened displayed a strong preference for (R) enantiomer in both reactions. A cluster 3 enzyme, PgOYE11 from *Pseudomonas sp.* GM48, and a cluster 16 enzyme, HmOYE6 from *Hypsizygus marmoreus*, both converted a racemic mix of **2p,** suggesting these OYEs are capable of the ‘flipped’ (rotated vs the axis of the carbonyls) or ‘reverse’ (opposite coordinated carbonyl) binding orientations when compared to other OYEs (Fig. 6B).^44^

**Figure 6:**
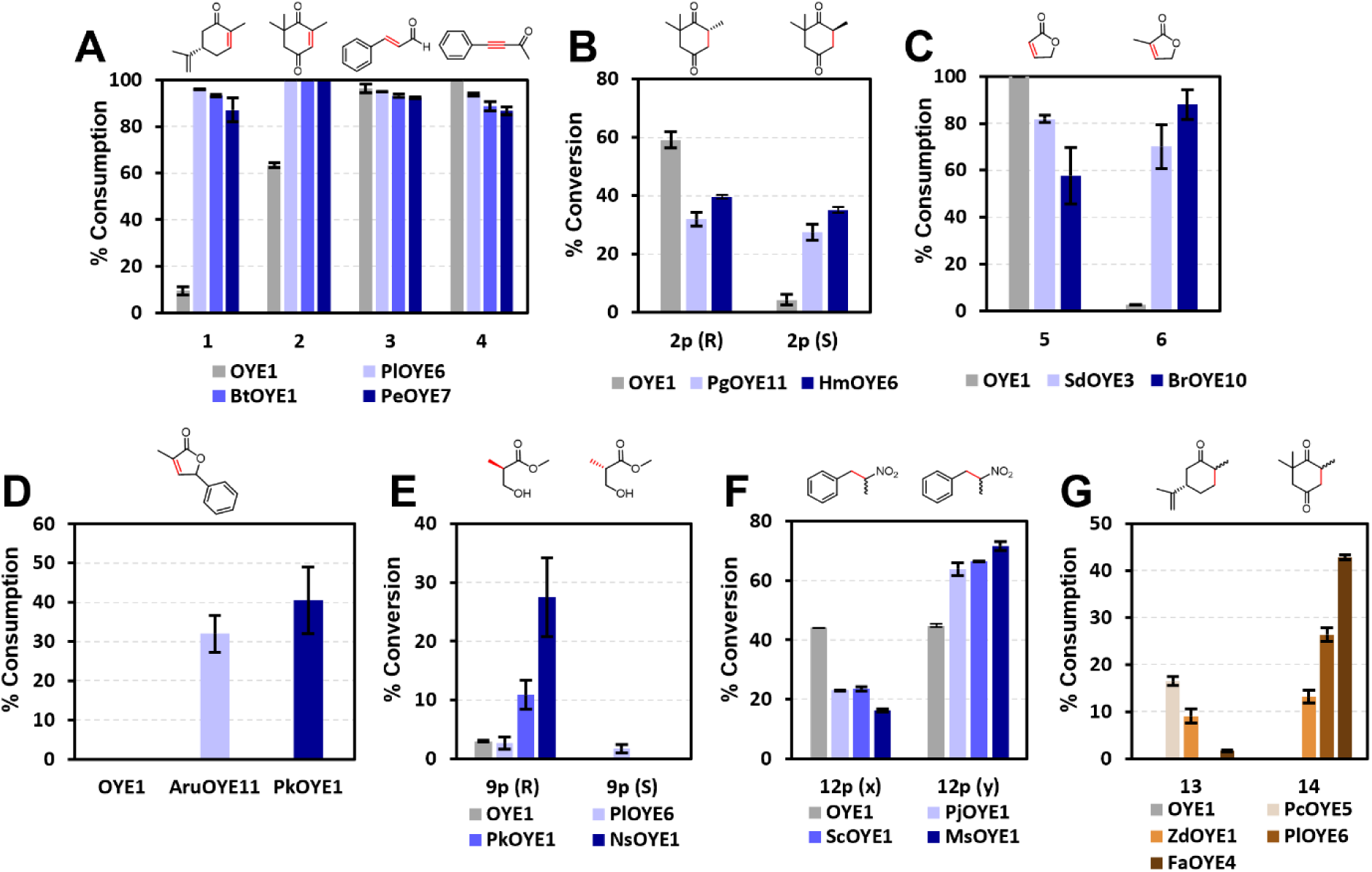
Activity of potential biocatalysts. The unique catalytic activity of potential biocatalysts is shown above. Colors represent the type of activity with blue for ene reduction and gold for desaturation. The substrate or product is shown above the graph.

Substrates **5** and **6** in mix II only differ by a methyl group. However, OYE1 heavily prefers **5** over **6** converting 100% vs <10%. Improved conversion of **6** was detected from two enzymes, SdOYE3 and BrOYE10 from *Sphingomonas dokdonensis* and *Bradyrhizobium sp*. R5 respectively (Fig. 6C). For substrate **8**, no conversion was detected for characterized enzymes. An OYE from cluster 11, PkOYE1 from an unidentified proteobacterium, and AruOYE11 from *Aspergillus ruber* and cluster 9, were the only enzymes to display activity for **8** (Fig. 6D). This bulky substrate likely indicates that their respective active sites may be slightly larger than other known OYEs. Interestingly, PkOYE1 does not reduce **5** but does convert the methylated **6,** showing a similar preference as the other cluster 11 enzyme, SdOYE3.

In mix III, **9** remains a challenging substrate for characterized OYEs. PkOYE1 and the cluster 1 enzyme NsOYE1, from *Neptuniibacter sp.,* convert 6- and 9-fold more of **9** when compared to OYE1 and other characterized enzymes (Fig. 6E). It is intriguing that PkOYE1 converts the bulkiest substrate (**8**) and one of the smallest substrates (**9**) at relatively high levels of conversion. Hailing from an uncharacterized cluster with no solved structures, it is unclear what the active site geometry of this enzyme is. Beyond improved conversion, PlOYE6 produced a racemic mix, whereas OYE1 and other enzymes heavily prefer the (R)-enantiomer. This could also prove to be the starting point for future mutational studies to produce the (S)-enantiomer. Lastly, alkene reduction of **12** results in a racemic mixture for the majority of enzymes except for cluster 1 enzymes: ScOYE1, MsOYE1, and PjOYE1 from *Stenotrophomonas chelatiphaga*, *Methylophaga sp*., and an unidentified proteobacterium, respectively (Fig. 6F). These enzymes all preferred the same enantiomer over the other, but the identity of the favored enantiomer could not be resolved. The methyl group at the α position of **12** creates the possibility of a stereocenter after reduction. Therefore, these 3 enzymes must have a preferred binding orientation for the substrate, likely a steric interaction with the methyl group.

While most of the desaturase activity for **13** and **14** was detected at low conversion levels, several enzymes displayed considerable oxidative capabilities (>10%). In particular, the two cluster 4 OYEs, PlOYE6 mentioned earlier and FaOYE4 from *Flavobacterium sp.* ABG, converted >25% of **14** within 24 hours with the latter enzyme reducing >40%. (Fig. 6G) For substrate **13,** the highest activity was detected for PcOYE5, a cluster 8 enzyme from *Pseudomonas chloritidismutans*, which has >15% conversion but could not convert **14** (Fig 6G). Several OYEs displayed substrate promiscuity for the desaturase reaction, such as ZdOYE1, a cluster 1 enzyme from an unclassified marine bacterium, that converted both **13** and **14** >10% (Fig 6G).

### Redox Potential of Desaturating OYEs

Few studies have explored the reverse, oxidative chemistry of OYEs.^37^ Massey et al. demonstrated that OYE1, an enzyme with no native desaturase activity, could be converted into a desaturase enzyme by increasing its redox potential via incorporation of artificial flavins.^38^ This suggests that the redox potential of an OYE may be critical to desaturase activity. The redox potentials of four OYEs were examined: PlOYE6, FaOYE4, PcOYE5, and ZdOYE1.^45^ OYE1 exhibited a redox potential midpoint of −227 mV, in agreeance with previous studies.^46, 47^ Interestingly a correlation between redox potential and desaturase activity was not observed: two OYEs, PcOYE5, and ZdOYE1, exhibited a more positive (+∼25 mV) redox potential than OYE1, but PlOYE6 was more negative (−20 mV) (Table 1). Compared to the artificial flavins used in previous studies (increase of 160 mV), the differences in redox potential between the enzymes with activity vs OYE1 are only slight.^38^ Furthermore, it has been demonstrated that raising the redox potential removed the ene reductase capabilities of OYEs. If an increased redox potential was the main factor in the observed desaturase activity, then the enzymes should be less capable of ene reduction. The opposite trend occurs as the enzymes with the highest desaturase activity coincidentally have the highest ene reductase activity (PlOYE6 and FaOYE4) for select substrates. Therefore, redox potential may influence the desaturase activity of certain OYEs, but other factors must also contribute, especially for the selected OYEs.

**Table 1:**
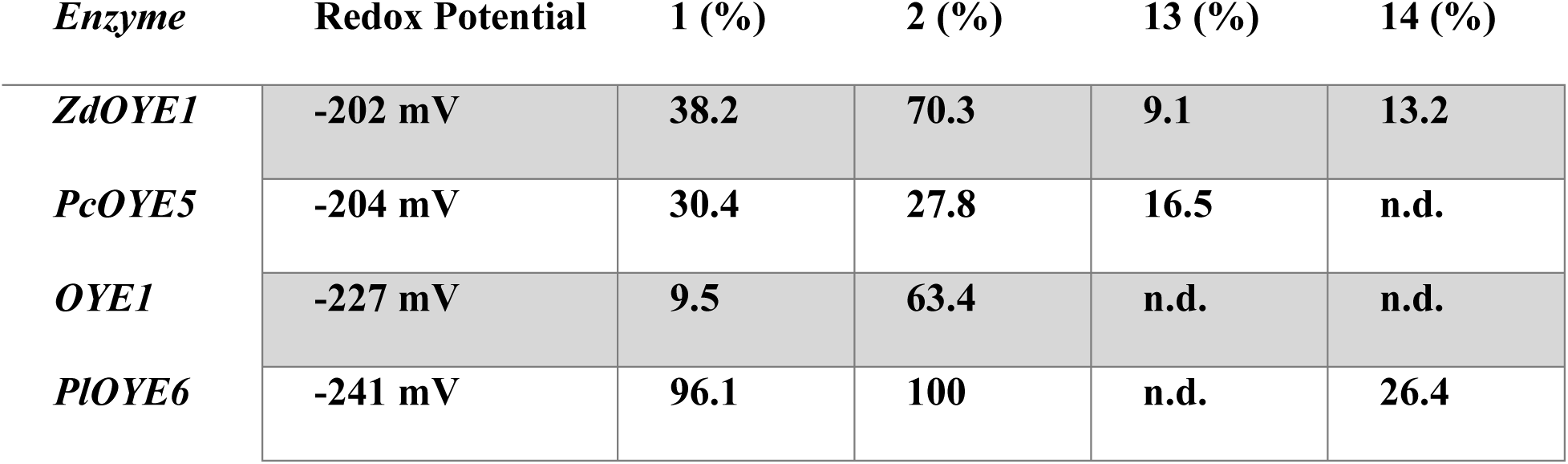
Redox potentials vs substrate consumption for selected OYEs. The desaturase OYEs are organized in decreasing value of redox potential. The percent consumption of mix 1 and mix 4 substrates are included to show ene reductase and desaturase of the enzymes.

## Conclusions

Here, a combination of bioinformatic and biochemical techniques was utilized to systematically assess the biocatalytic potential of the large OYE family. SSN analysis identified several previously unknown areas within the sequence space, including unique OYE subclasses such as cluster 8. Overall, 119 novel, diverse representative sequences were screened with 60% showing activity - representing a >50% increase in the total number of characterized OYEs. Surprisingly, desaturase activity was observed throughout the enzyme family which was previously concluded to only occur at elevated temperatures or cofactor manipulation. This significantly expands the enzymatic versatility of OYEs. Despite previous studies, the observed desaturase activity does not appear to be heavily influenced by an enzyme’s redox potential as no correlation was observed between the two. Thus, these findings suggest that desaturase activity in OYEs must be dependent on other factors.

In total, 14 novel enzymes were identified as potential biocatalysts. Of note, PlOYE6 is an excellent ene reductases converting all substrates of mix I, while also exhibiting significant desaturase activity specifically for **14** and the highest activity for **9p** (S). Two enzymes, PkOYE1 and NsOYE1, showed enhanced conversion (6- and 9-fold) for the challenging substrate **9**. Additionally, PkOYE1 and AruOYE11 convert the large bulky **8** which OYE1 and the other characterized enzymes did not. Enzymes with unusual enantioselectivity and improved catalysis were identified in every mix and are promising starting points as possible industrial biocatalysts. To harness the full potential of biocatalysis for sustainable chemical production, new enzymes must be continuously discovered. In the postgenomic era, with an overwhelming amount of sequence info, the emphasis has shifted from identifying new enzymes to selecting which to characterize. This combined strategy of bioinformatics and synthetic biology has revealed a wealth of knowledge concerning OYEs while also detecting possible biocatalysts candidates. As a platform for protein family screening, a similar approach could be applied to explore the biocatalytic potential of other large, diverse enzyme families in order to meet the need for biocatalysis.

## Author Contributions

The manuscript was written through the contributions of all authors.

## Acknowledgments

We thank Dr. Melanie Hall (University of Graz) for providing a sample of 3-methyl-5- phenyl-5H-furan-2-one (**8**). Helpful comments and suggestions from members of the Atlanta Flavin Group and the Lutz lab are acknowledged.

## Abbreviations

OYE: old yellow enzyme
SSN: sequence similarity network
IVTT: in vitro transcription/translation system
FMN: Flavin mononucleotide
FAD: Flavin adenine dinucleotide

## Materials and Methods

### Generation of SSNs

Sequence similarity networks were generated using the EFI-EST^48^. In brief, 70,366 protein sequences associated with the PFAM designation PF00724 ‘OYE-Like’^49^ were collected, and an edge-threshold of 10^-85^ was selected to separate known enzyme classes (Classes I, II, and 2-enoate reductases) into distinct clusters.

### Generation of Phylogenetic Trees

A comprehensive phylogenetic tree for the OYE family was constructed using a proportional representative selection of from the 20 major clusters and including all characterized sequences (inclusive of this study). This resulted in a set of 470 sequences. The enzymes were aligned using PROMALS3D.^50^ In the event of multidomain proteins, only the OYE-like domain was used. The resulting clustal alignment was used to generate a phylogenetic tree with IQTree using the WAG substitution model and freerate heterogeneity.^51^ The tree was visualized using iTOL.^52^

### Determination of the Solubility of Selected OYEs

OYE sequences were expressed in BL21 cells in 5 mL 2xYT media with appropriate antibiotic. Initial growth was incubated at 37°C until an OD600 of 0.6 was observed. Then the cells were induced using Isopropyl β-D-1-thiogalactopyranoside (IPTG) to a final concentration of 0.3 mM. The cells expressed at 20°C overnight and the next day were spun down at 4000 g at 3 min to pellets which were chemically lysed with BugBuster (EMD). The solutions were rocked for 20 min to fully lyse the cells. Then the samples were centrifuged for 10 min at 8000 g. The supernatant (soluble) was removed, and the resulting pellet (insoluble) was resuspended with DI water. Both fractions were loaded onto an SDS-PAGE gel for solubility analysis.

### Expression of OYE Sequences using Modified PURExpress

Selected novel OYE sequences were codon optimized for *E. coli* expression then synthesized/placed into pET28a plasmids by the DOE Joint Genome Institute. For in vitro expression of selected sequences, the PURExpress kit from New England Biolabs was used for rapid production of the enzymes. Here in vitro expression reactions were performed in duplicates with the following components: PURExpress Solution A, Solution B, 200 ng of Template DNA, 100 μM FMN and FAD, 5 μM of GroEL, 10 μM GroES, 20 U Murine RNAse Inhibitor, and RNAse Free Water. FMN and FAD were added to fulfill the potential cofactor requirements of the novel OYE sequences. GroES and GroEL were added to improve proper protein folding of the target enzyme. Purification and quantification of both GroES and GroEL were based on literature.^53^ In addition to the novel OYE sequences, a positive control of OYE1 in pET14b and negative control of dihydrofolate were expressed with each batch of in vitro expression. Reactions were incubated at 37 C for 2.5 hours then quenched on ice before aliquots were added into an activity assay.

### Activity Assay Conditions for Novel OYE Sequences

To determine the ene reductase and desaturase activity of selected OYE sequences, in vitro expressed protein was aliquoted into reaction mixtures and allowed to react for 24 hours. Substrates were purchased from Sigma Aldrich except for **8** which was a gift from Melanie Hall at the University of Graz. For mixes I-III (ene reductase), reaction mixtures contained 2 U of glucose dehydrogenase, 100 mM glucose, 200 μM of NADH and NADPH and 250 μM (1 mM for mix III) of each substrate in a buffer of 50 mM Tris HCL pH 7.5. Glucose dehydrogenase and glucose were added to recycle NAD(P)H to ensure sufficient reducing equivalents. Purification of this enzyme was performed using methods from literature.^54^ Reactions also occurred under anaerobic conditions to prevent competition with oxygen. For mix IV (desaturase), reaction mixtures only contained 20 mM pyrophosphate buffer ph 8.5 with 250 μM of each substrate under aerobic conditions. Reaction conditions for desaturase activity were adapted from literature.^38^ Each reaction was performed in duplicate and after 24 hours were quenched using ethyl acetate. In addition, substrates and products were extracted twice using ethyl acetate containing 500 μM cyclohexane (internal standard) then placed onto GC/MS for analysis.

### GC/MS Parameters

GC/MS quantification was performed using Shimadzu QP2010 SE instrument using an FID detector with helium as a carrier gas. A chiral CycloSil-B column was used to separate the compounds. Mixture GC/MS protocols were developed and are as follows: *Mix I/IV*: Injection temperature 250 °C, oven 110 °C with ramp to 150 °C at 2 °C/min, hold for 5 min. *Mix II*: Injection temperature 220 °C, oven 80 °C hold for 1 min then ramp to 100 °C at 1 °C/min hold for 1 min then ramp to 185 °C at 2.5 °C/min. *Mix III*: Injection temperature 210 °C, oven 75 °C hold for 5 min then ramp to 100 °C at 2 °C/min, ramp to 200 °C at 10 °C/min hold for 5 min.

### Expression and Purification of Selected OYE Sequences

Plasmids were transformed into chemically competent E. coli BL21 (DE3) cells and spread onto kanamycin (30 mg/mL) agar plates and allowed to grow overnight at 37°C. Colonies were picked and 5 mL LB with kanamycin were inoculated overnight. The next day, 1 mL of the overnight culture was used to inoculate 250 mL of 2xYT medium (with kanamycin) for expression. The culture was allowed to grow until an OD600 of 0.55-0.7 was reached. At the desired OD, the expression cultures were induced with 0.3 mM IPTG and expressed for 18-22 hours for 20°C. After expression cells were poured into tubes and spun at 4000 g for 20 minutes. The supernatant was decanted, and tubes were placed into freezer (−20°C) to aid with lysing.

Frozen cell pellets were thawed on ice until liquid then resuspended in buffer 1 (40 mM Tris-HCl pH 8.0, 10 mM NaCl) with 2.5 μL/g of DNase I and 50 μL/g of protease inhibitor cocktail. The solution was chilled on ice for 20 minutes and then sonicated for 3 minutes (10-sec pulse/20-sec off). The mixture was centrifuged for 30 minutes at 4000 g and 4°C. The supernatant was loaded onto a 5 mL HiTrap Q FastFlow anion-exchange column. The column was washed with buffer 1 for 2 column volumes (CV). Over the course of 15-20 CVs, a linear gradient from buffer 1 to 100% buffer 2 (40 mM Tris-HCl pH 8.0, 1 M NaCl) was used to elute the protein, identifying via the UV-vis absorption of the protein (280 nm) and flavin (450 nm). The protein was collected and concentrated using Millipore filter unit with 10 kDa MWCO. A size exclusion column (Superdex 200, 10/300 GL column) was used to finish purification. Fractions for both columns were analyzed via SDS-PAGE to ensure purity.

### Xanthine-Xanthine Oxidase Assay for OYE Redox Determination

In this assay, xanthine oxidase was used to slowly provide electrons for reduction of the enzyme and a reference dye while monitoring using a Cary UV-vis spectrometer. As OYE1 has a literature redox value of −227 mV, anthraquinone-2-sulfonate, −225 mV, was selected as the reference dye.^45, 47^ Other dyes were considered and utilized such as cresyl violet (−166 mV) and phenosafranine (−252) but were too far in redox potential for most OYEs to be properly measured. In an airtight vial, 20 μM of the target OYE, 20 μM methyl viologen, 400 μM xanthine, and 40 μM anthraquinone-2-sulfonate were added to 50 mM NaPO_4_ buffer at pH 7. In a glass bulb in the side of the cuvette, 100 nM xanthine oxidase was added. The cuvette was subjected to 10 rounds of vacuum and Ar to fully deoxygenate the solution. An initial scan was taken before the reaction, then the solution was tipped to add the xanthine oxidase, and UV-vis spectra were collected. The concentration of reduced and oxidized species were determined using absorbance with 331 nm for anthraquinone-2-sulfonate and 460 nm for selected OYEs. Redox potential was calculated using a derived Nernst Equation (1) and explained in further detail in the literature.^45^

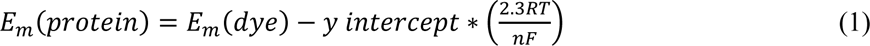

F is the faradaic constant and n the number of electrons transferred which is 2. The y-intercept is from the plot of log oxidized/reduced protein vs dye.

