## Supplemental for "The Hidden Biocatalytic Potential of the Old Yellow Enzyme Family"

^1^Department of Chemistry, Emory University,
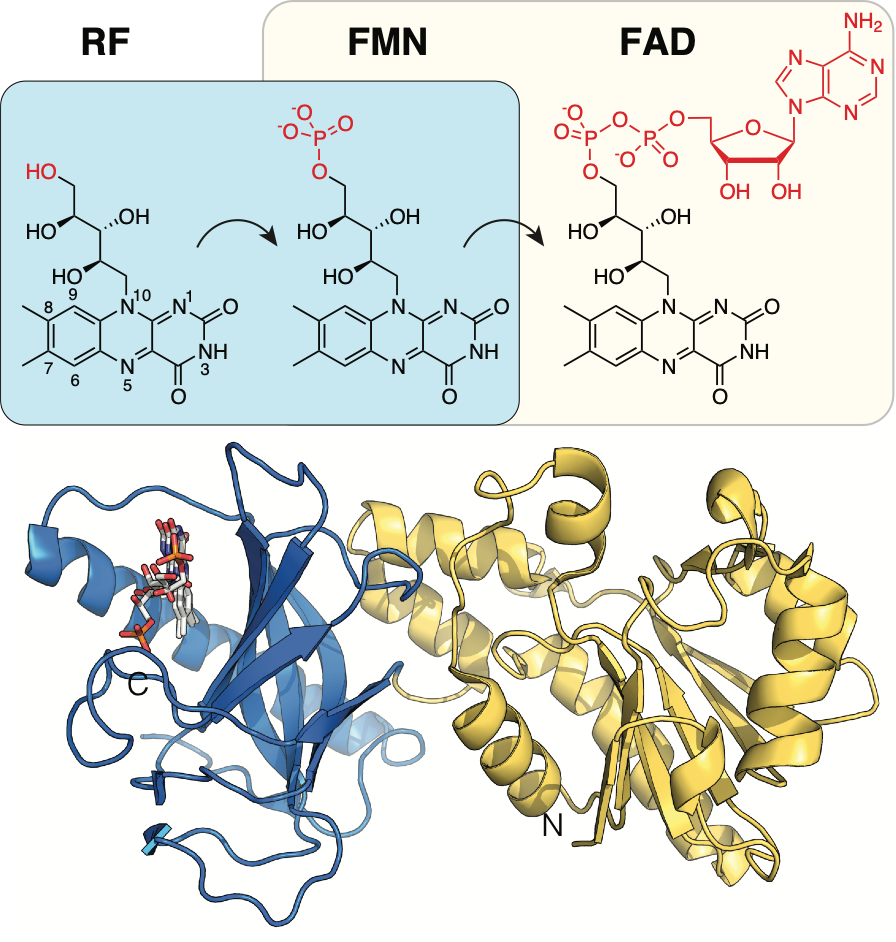
 Atlanta, Georgia 30084, United States

^2^Michael Smith Laboratories, University of British Columbia, Vancouver, BC, Canada

Table S1 Summary of OYE Library.

| UniProt | Generation | Cluster |
| --- | --- | --- |
| B9T8J4 | ▲81 3D7 | 3 |
| M5B1C9 | 2E12 | 61 |
| V7I5A8 | 2H12 | 15 |
| W7ZMG5 | 2H7 | 8 |
| A0YFJ6 | 3A3 | 11 |
| B9Y7J4 | 3A4 | 20 |
| Q9JN79 | 3A5 | 4 |
| Q9WYQ9 | 3A7 | 14 |
| R7SZ38 | 3A8 | 22 |
| S6WB48 | 3B1 | 2 |
| Q24RN8 | 3B2 | 63 |
| P54524 | 3B4 | 6 |
| C0ZVN3 | 3B6 | 7 |
| A0A017SC09 | 3B7 | 9 |
| V4RWU7 | 3C4 | 8 |
| Q4AA30 | 3C5 | 13 |
| A8FEW4 | 3D1 | 3 |
| Q4WZ70 | 3D4 | 1 |
| W0DA85 | 3E1 | 3 |
| K6TQP5 | 3E2 | 25 |
| P77258 | 3F3 | 1 |
| M2Z4K1 | 3F7 | 26 |
| C3MNJ2 | 3F8 | 2 |
| K0B444 | 3G1 | 34 |
| Q8PUE0 | 3G3 | 10 |
| Q2TJB8 | 3G4 | 1 |
| S5YUA6 | 3G5 | 29 |
| J7L5I6 | 3G6 | 5 |
| C0QFC4 | 3H3 | 10 |
| Q02899 | 3H5 | 1 |
| J3P9F5 | 3H6 | 3 |
| E3HDQ4 | A1 | 21 |
| E9E863 | A10 | 1 |
| A0A1Q8DL33 | A11 | 17 |
| A0A1W0XDZ4 | A12 | 1 |
| A0A1H1DXC0 | A2 | 3 |
| A0A1H3H521 | A3 | 3 |
| A0A1Q3H1G5 | A4 | 1 |
| A0A104JBL3 | A5 | 1 |
| U9YMT6 | A6 | 6 |
| M2VAD5 | A7 | 3 |
| A0A1C4MT72 | A8 | 1 |
| A0A2D1JR17 | A9 | 6 |
| A0A062X679 | B1 | 3 |
| A0A1G6QTQ7 | B10 | 3 |
| A0A1Y1W9M1 | B11 | 3 |
| B8LTL5 | B12 | 3 |
| A5VBP0 | B2 | 1 |
| A0A0R0D6E4 | B3 | 1 |
| A0A1X6MY35 | B4 | 1 |
| C7ZM49 | B5 | 1 |
| A0A2G0VWT1 | B6 | 8 |
| A0A2C2U3H6 | B7 | 6 |
| A0A1Z9EHY4 | B8 | 3 |
| A0A0U1LWY3 | B9 | 1 |
| R4K9R9 | C1 | 3 |
| G4HUZ7 | C10 | 7 |
| Q6CUW9 | C11 | 3 |
| A0A1S9RTM4 | C12 | 3 |
| A0A0P7GE52 | C2 | 1 |
| A0A0M4QXT8 | C3 | 3 |
| A0A1Y1YHM2 | C4 | 1 |
| A0A0Q7WCQ9 | C5 | 1 |
| A0A0B7GCC5 | C6 | 1 |
| A0A0F9NP68 | C7 | 10 |
| D8M5A9 | C8 | 3 |
| W8NW54 | C9 | 1 |
| A0A0H4KFX4 | D1 | 3 |
| A0A226DQD1 | D10 | 3 |
| A0A0B7N6T2 | D11 | 3 |
| V5TJE2 | D12 | 18 |
| A0A2E0KBB9 | D2 | 1 |
| A0A2G2QFR0 | D3 | 1 |
| A0A245ZID8 | D4 | 11 |
| A0A087B547 | D5 | 3 |
| A0A0D6I6F4 | D6 | 4 |
| A0A137SCF2 | D7 | 1 |
| A0A1D1W904 | D8 | 3 |
| J3GED7 | D9 | 3 |
| A0A1Q4W807 | E1 | 3 |
| K5Y813 | E10 | 3 |
| A0A1H1PA63 | E11 | 7 |
| Q3JA76 | E12 | 23 |
| A0A1Q5C6Q4 | E2 | 3 |
| W9ATX1 | E3 | 3 |
| A0A239MCT0 | E4 | 1 |
| A0A098SST3 | E5 | 4 |
| A0A0C2I951 | E6 | 1 |
| U6SIA5 | E7 | 6 |
| J3EXQ1 | E8 | 8 |
| A0A1A0W0R3 | E9 | 4 |
| A0A166JDD2 | F1 | 1 |
| A0A2A3FEX1 | F10 | 7 |
| L7ITH5 | F11 | 9 |
| Q23090 | F12 | 9 |
| A0A0D5AF70 | F2 | 3 |
| A0A2D5KJ34 | F3 | 11 |
| A0A1I6Q9M7 | F4 | 1 |
| A0A1X2BF87 | F5 | 1 |
| A0A2A9JE68 | F6 | 3 |
| K3WFC3 | F7 | 1 |
| A0A100YAA1 | F8 | 1 |
| D0NQX8 | F9 | 1 |
| A0A014MP39 | G1 | 1 |
| E3LUM4 | G10 | 9 |
| G3JMK4 | G11 | 5 |
| A0A136H5V4 | G2 | 1 |
| A0A0E3SPA7 | G3 | 1 |
| B5XWM6 | G4 | 1 |
| Q87XC7 | G5 | 1 |
| A0A161P8U3 | G6 | 4 |
| A0A0S3C0K6 | G7 | 12 |
| A0A151VNU3 | G8 | 16 |
| A0A0A2JDP3 | G9 | 24 |
| A0A1I5G981 | H1 | 3 |
| A0A1H1V287 | H10 | 5 |
| A0A261Y278 | H11 | 9 |
| A0A1R0ZM98 | H2 | 1 |
| A0A1M7EKV3 | H3 | 1 |
| P71278 | H4 | 1 |
| A0A0J0YHR8 | H5 | 4 |
| A0A1M5T1E8 | H6 | 10 |
| A0A061QIQ1 | H7 | 1 |
| A0A2G4DT26 | H8 | 3 |
| A0A0R3L3J0 | H9 | 3 |
